# Direct observation of feeding behavior in the kleptoplastic sea slug *Plakobranchus ocellatus* type black (Gastropoda, Sacoglossa)*

**DOI:** 10.64898/2026.01.23.701294

**Authors:** Shinichi Inagaki, Taro Maeda

## Abstract

The feeding ecology of sacoglossans, an exceptionally stenophagous group among marine animals, has been extensively studied because these organisms exhibit kleptoplasty—a phenomenon in which chloroplasts derived from ingested algae are sequestered and functionally maintained within animal cells. The genus *Plakobranchus* is notable for its exceptional capacity to retain photosynthetically active chloroplasts for prolonged periods. Nevertheless, direct observations of its feeding behavior have remained limited, largely due to subsequent taxonomic revisions and its cryptic, benthic lifestyle. In the present study, we focus on *Plakobranchus ocellatus* type black (PoB). Using high-resolution macro videography, we obtained, for the first time, video recordings that directly document the feeding behavior of this lineage. Under illuminated conditions, we demonstrate active feeding on the fan-shaped fronds of udoteacean algae, clarify which algal life-history stages serve as food, and identify *Ventalia* sp. 4 as a previously unrecognized dietary component. We further describe a characteristic inverted “headstand” posture adopted during feeding and discuss this behavior in relation to the species’ benthic and semi-burrowing lifestyle, proposing that it may partly explain the historical rarity of feeding observations in the field. The confirmation of feeding behavior represents a critical step toward establishing a stable laboratory culture system essential for experimental studies of kleptoplasty.

## Main Body

Kleptoplasty—the retention of functional algal chloroplasts in animal cells—represents one of the remarkable examples of inter-kingdom organelle acquisition (Miyagishima, 2023).

Among sacoglossan sea slugs, the genus *Plakobranchus* van Hasselt, 1824 exhibits particularly long-term photosynthetic activity, retaining functional plastids for several months (Evertsen *et al*., 2007). However, despite extensive research on chloroplast physiology, the feeding behavior of *Plakobranchus* remains poorly understood, largely due to its cryptic lifestyle and taxonomic complexity.

The genus *Plakobranchus* is among the earliest described taxa within the order Sacoglossa van Ihering (1876) (Van Hasselt, 1824; Jensen, 1992; Meyers-Muñoz *et al*., 2016). Although more than a dozen nominal species were historically described, their morphological similarity led to synonymization under *P. ocellatus sensu lato* (Jensen, 1992; Meyers-Muñoz *et al*., 2016). Molecular analyses, however, have since revealed five to eight genetically distinct *P. ocellatus s*.*l*. lineages (operationally defined as “types”), with *P. papua* Meyers-Muñoz *et al*., 2016, *P. noctisstellatus* Mehrotra, Caballer, C. M. Scott, Sp. Arnold, Monchanin and Chavanich, 2020, and *P*. cf. *ianthobaptus* Gould, 1852 now recognized as separate species (Trowbridge *et al*., 2011; Krug *et al*., 2013; Meyers-Muñoz *et al*., 2016; Mehrotra *et al*., 2020). This taxonomic complexity has resulted in fragmented physiological data, often mixing observations from multiple lineages.

Feeding behavior is similarly poorly known: individuals are usually found resting on sandy substrates, rarely on algae, and only three feeding events—on *Rhipidosiphon javensis* and *Chlorodesmis hildebrandtii*—were ever documented (Kawaguti, 1941; Dunlap, 1975; Jensen, 1992). Furthermore, it remains unclear which lineage those researched *Plakobranchus* individuals belonged to. Subsequent molecular studies of *Plakobranchus* identified multiple potential algal sources, including *Rhipidosiphon* and *Chlorodesmis* species, but no direct observations confirmed their ingestion (Wägele *et al*., 2011; Maeda *et al*., 2012; Christa *et al*., 2012; Wade & Sherwood, 2017, 2018).

We focused on *P. ocellatus* type black (PoB), a molecularly defined lineage for which both genomic data and kleptoplast barcoding exist (Maeda *et al*., 2012, 2021; Krug *et al*., 2013) (Figures 1A–C). Previous analyses identified *R. lewmanomontiae* as one of the main kleptoplast sources (Figure 1D), and stable isotope data suggest active feeding, but behavior had not been recorded (Maeda *et al*., 2012).

**Figure 1.**
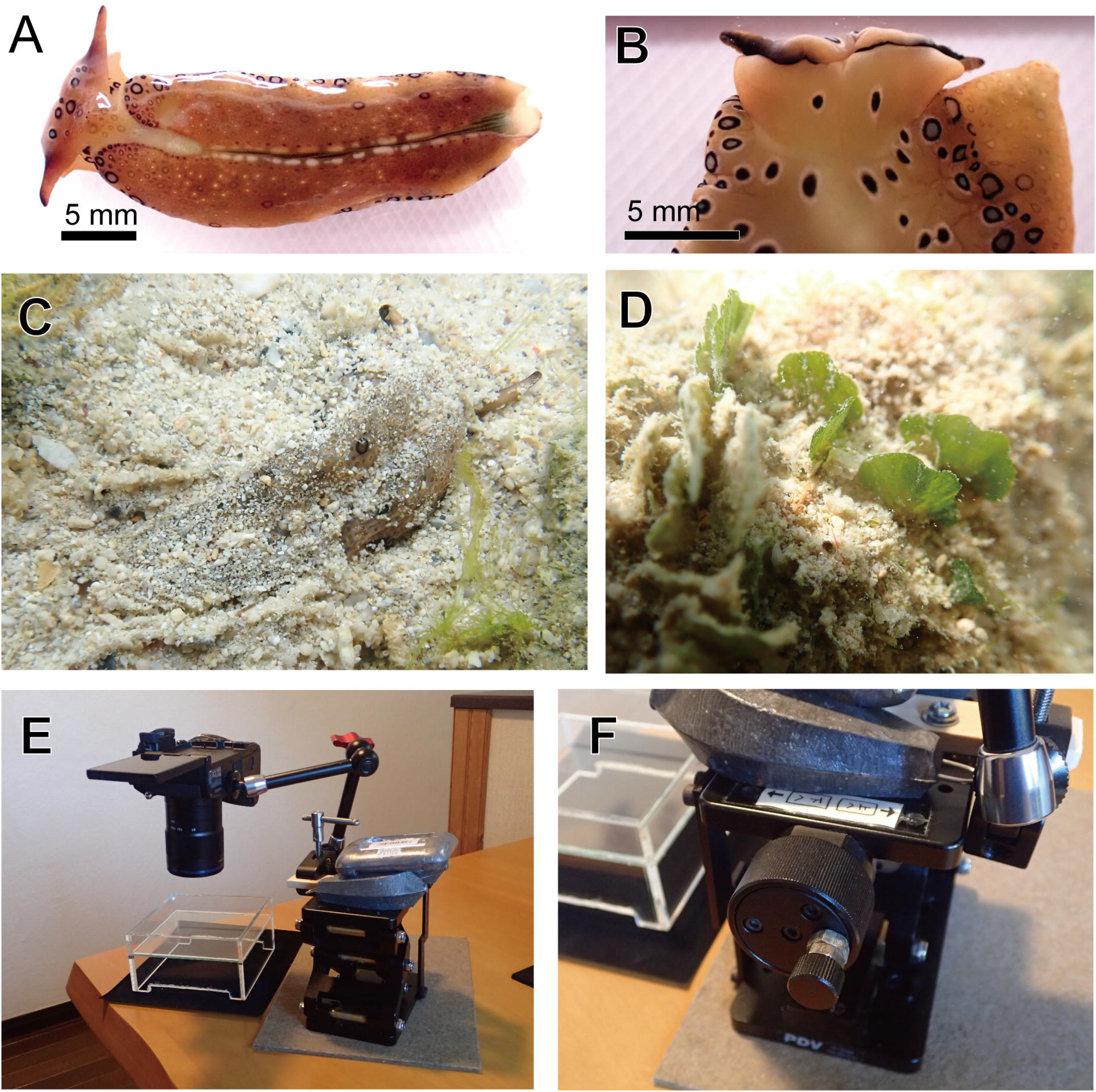
Overview of *Plakobranchus ocellatus* type black (PoB) examined in this study. **A**. Dorsal view of PoB. **B**. Ventral view of the head part. **C**. PoB partially buried in sand in its natural habitat (body length ∼3 cm). **D**. Field photograph of the algal food source, *Rhipidosiphon* cf. *lewmanomontiae*; each fan-shaped frond measures approximately 5 mm in diameter. **E**. Camera setup for recording feeding behavior. **F**. Platform design used for macro photography.

Samples of PoB and their potential algal food sources were collected by snorkeling and manual sampling in intertidal zones of Japan (Table 1). Molecular phylogenetic analyses based on chloroplast *rbcL* sequences were conducted to identify the collected algal specimens, following the taxonomic framework of Lagourgue & Payri, 2022.

**Table 1.**
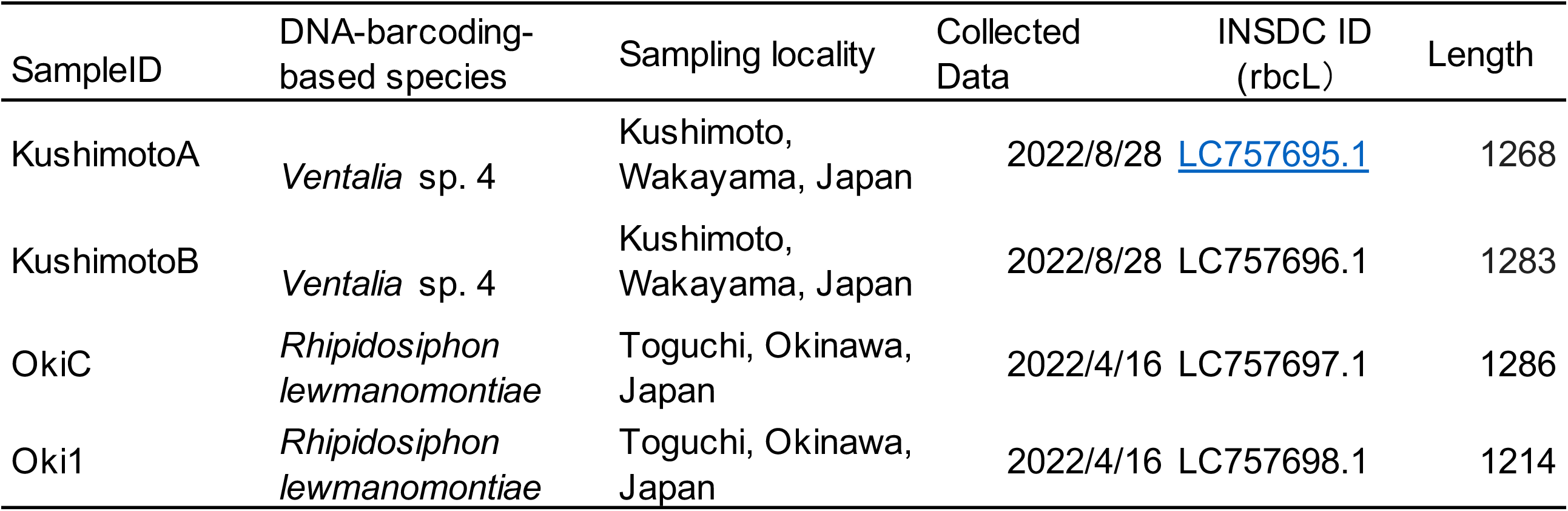
Collection data and INSDC accession information for specimens used in this study.

Algal DNA was extracted from either the grazed thalli or nearby individuals using the DNeasy Plant Mini Kit (Qiagen), following the manufacturer’s instructions. The *rbcL* region was amplified with newly designed Rhipidosiphon-specific primers (rbc1_Rhip: 5 ′ - CCACAAACAGAAACAAAAGC-3′; U3-2_Rhip: 5′-TCTTTCCATACAACACAAAGC-3′) using KOD One PCR Master Mix (Toyobo, Osaka, Japan). Amplicons were purified with NucleoSpin® Gel and PCR Clean-up Kit (Takara Bio, Shiga, Japan) and Sanger-sequenced by Eurofins Genomics (Ebersberg, Germany). Sequences were aligned with MAFFT v7.526, trimmed with trimAl v1.5, and subjected to maximum-likelihood phylogenetic analysis using IQ-TREE v2.3.6. All sequences have been deposited in INSDC (Table 1).

Animals were maintained in aquaria at 25 °C under starvation for one month. Feeding behavior was recorded using an OM-D E-M1 Mark II camera (OM Digital Solutions, Tokyo, Japan) equipped with a LAOWA 50 mm F2.8 2× Ultra Macro APO lens (Venus Optics, Hefei, China), in a 12 × 2 × 12 cm glass tank (Figure 1E, F). Starved PoB individuals were introduced to aquaria containing a single algal species, and their feeding behavior was directly observed and recorded.

DNA barcoding identified the algal specimens from Okinawa and Wakayama as *R. lewmanomontiae* and *Ventalia* sp. 4 (sensu Lagourgue & Payri, 2022), respectively. Partial *rbcL* sequences (1,214–1,286 bp) obtained from four thalli (two per locality) showed 99.7–99.8% identity with published reference sequences of these taxa (HQ871689, MT339698). Maximum-likelihood phylogenetic analysis based on *rbcL* confirmed these identifications, with both clades receiving high bootstrap support (> 99%) (Figure 2A). Morphologically, both algae exhibited fan-shaped fronds typical of inconspicuous *Rhipidosiphon* and *Ventalia* species, and no morphological differences beyond individual or environmental variation were detected as mentioned in Lagourgue & Payri, 2022 (Figure 2B). The present finding extends the known range of *Ventalia* sp. 4 northward to Wakayama, representing its first record outside Southeast Asia (Lagourgue & Payri, 2022).

**Figure 2.**
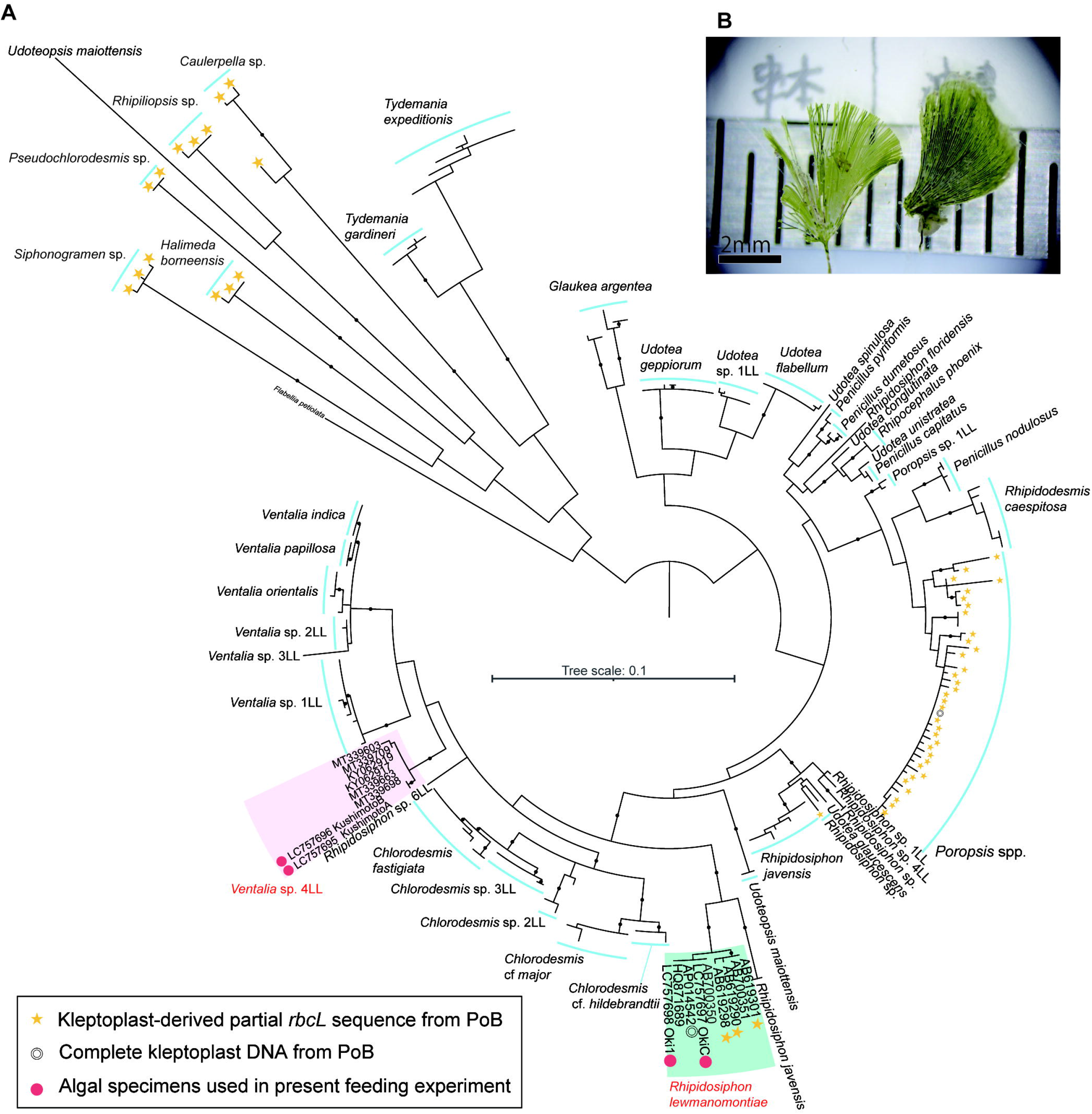
Identification of algal species used in feeding experiments. **A**. Maximum-likelihood phylogenetic tree constructed from *rbcL* sequences (715 positions) of related Bryopsidalean algae. Nodes supported by bootstrap values ≥99 are indicated by black circles. The complete phylogenetic tree is provided in Supplementary Figure 1. **B**. Algal thalli used for analysis: specimens from Wakayama (left) and Okinawa (right).

Feeding behavior was consistent across both algal species (Figures 3A–N, Supplementary Videos 1 and 2). Starved PoB began feeding within minutes after contact with *R. lewmanomontiae* or *Ventalia* sp. 4. The slugs approached algal thalli, probed with their lips, and grasped the fronds using a midline oral groove to draw tissue toward the mouth (Figures 3A, B, G, H, K–M). Feeding occurred on the fan–shaped frond rather than on the stipes of thalli.

**Figure 3.**
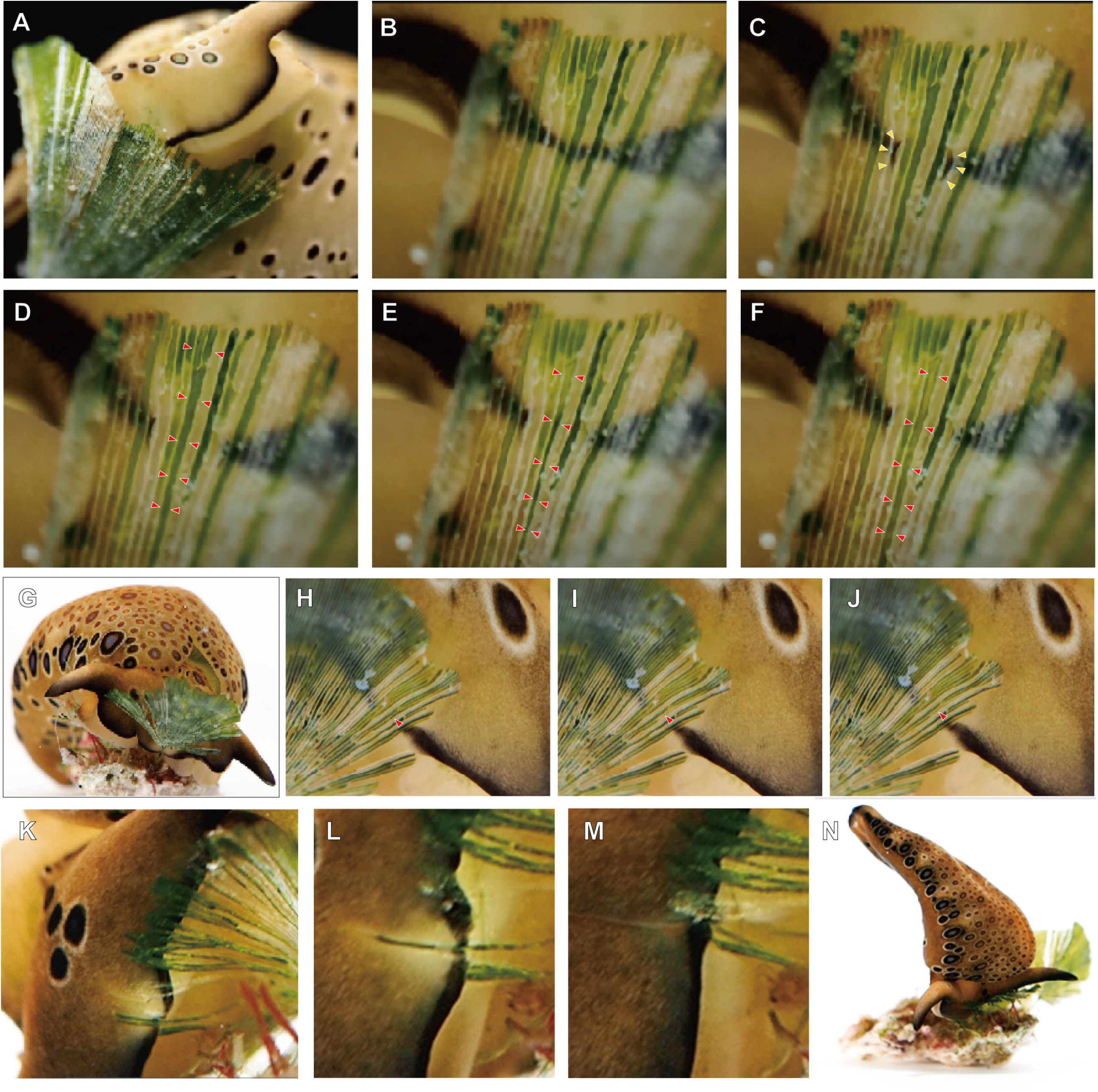
Feeding behavior of *P. ocellatus* type black (PoB). **A–F**. PoB feeding on *R. lewmanomontiae*. **G–H**. PoB feeding on *Ventalia* sp. 4. **A**. PoB initiating feeding at the distal tip of a fan-shaped frond. **B–F**. Sequential close-up images of the lip region during feeding. **B**. Lips pressed firmly against the algal surface. **C**. Opening of the oral apparatus (yellow arrow: black marginal line of the oral opening). **D–F**. Red arrows indicate algal cells being ingested. **D**. Algal cells prior to ingestion. **E**. Cells deformed by suction. **F**. Cells emptied of their contents after ingestion. **G**. PoB initiating feeding from the central region of a fan-shaped frond of *Ventalia* sp. 4. **H–J**. Red arrows indicate algal cells undergoing ingestion. **H**. Algal cells before ingestion. **I**. Cells after suction. **J**. Cells with contents expelled following regurgitation. **K–M**. PoB grasping and ingesting the fibrous elements of a disintegrating frond with its lips. **N**. Individual adopting an inverted posture during feeding. See Supplementary Videos 1 and 2.

Observed individuals began feeding either from the distal tip (Figure 3A) or the central portion of the fan-shaped frond (Figure 3B). PoB fed by wrapping its lips around individual algal filaments (Figures 3K–M), particularly at loosely connected sites, whereas in more robust regions it pressed its head firmly against the algal surface (Figures 3B, H). During feeding, the distal portion of the buccal apparatus protruded forward from the lips and was visible through the algal tissue, bearing a distinct black ring-shaped spot at its tip (Figures 3B–C). PoB also exhibited regurgitative behavior, repeatedly expelling and re-ingesting partially digested algal cytoplasm (Figures 3H–J), a process that likely enhances enzymatic mixing and nutrient absorption (Jensen, 1981, 1997). In several cases, individuals pressed their heads against the algal surface while elevating the posterior body in a characteristic “headstand” posture (Figure 3N).

Our observations provide the first direct documentation of feeding in PoB and confirm and show the ingestion of *R. lewmanomontiae* and *Ventalia* sp. 4. The consumption of *R. lewmanomontiae* is consistent with previous DNA-barcoding results (Maeda *et al*., 2012), but molecular data alone could not reveal which life-history stage serves as food. The upright, fan-shaped frond of *R. lewmanomontiae* develops from an endolithic filamentous phase that bores into coral rubble (Drew & Abel, 1988; Vroom *et al*., 2001; Lagourgue & Payri, 2022). Previously, the absence of direct feeding observations had led to the hypothesis that PoB might graze on turf-forming microscopic filaments partially emerging from coral rubble (Maeda *et al*., 2012). However, our observations demonstrate that feeding occurs on macroscopically visible frond, not on filamentous stages nor holdfast (Figure 3). Feeding on holdfast structures was never observed, and the consumption of gametangial tissues remains uncertain because this developmental stage was not available during our experiments..

Feeding on *Ventalia* sp. 4, which was previously undetected in cloning-based DNA-barcoding (Maeda *et al*., 2012), demonstrates its potential inclusion in PoB’s natural diet. Although *Plakobranchus* is not distributed on Wakayama, with its northernmost confirmed range being Tanegashima Island in Japan (Kimoto, 2025), *Ventalia* sp. 4 is likely to be widely distributed along the Black Current system and is expected to overlap broadly with the distribution of PoB. Further surveys are warranted to assess the diversity of these fan-shaped algae and may uncover a broader natural spectrum of algal species serving as food for PoB. In the closely related species *P*. cf. *ianthobaptus* from Hawaii, the inferred breadth of algal food sources increased from 11 taxa (Wade & Sherwood, 2017), as detected by cloning-based DNA barcoding, to nearly 20 taxa based on multi-site Illumina sequencing surveys (Wade & Sherwood, 2018). Another study has argued that the duration of plastid retention in *P. ocellatus s. l*. can vary depending on the algal source (Christa *et al*., 2012). Thus, *Ventalia* sp. 4 may represent a minor food source that is difficult to detect using cloning-based DNA barcoding and/or supplies only short-lived plastids. Integrating molecular surveys with direct and controlled feeding experiments may further elucidate the breadth of algal associations in *Plakobranchus*.

Behaviorally, PoB perforates algal cell walls and sucks out cytoplasmic contents, like other sacoglossans. However, its characteristic “headstand” posture appears unique to this species. This stance may facilitate feeding on algae that are partially buried in sediment or attached to silt-covered substrates (see Figure 1D), consistent with PoB’s benthic, burrowing lifestyle. Pressing the head against the frond likely improves radular adhesion and efficiency of perforation, compensating for its limited ability to grasp thalli. On sandy substrates, PoB may gradually bury its head while following partially buried algal fronds via this “headstand” behavior, thereby gaining access to adjacent fronds arising from the same endolithic filaments within the sediment. Alternatively, this posture could simply reflect reduced foot mobility.

Although the species secretes adhesive mucus, it is unable to firmly grasp algal tissues, which may explain its inability to feed on detached thalli. The rarity of feeding observations in nature may therefore stem from this inconspicuous posture—if PoB typically feeds while partly buried or inverted against substrates, such behavior would be difficult to detect *in situ*. In addition, the headstand posture on exposed rock surfaces may easily collapse under water flow disturbances caused by researchers.

Feeding in *Plakobranchus* has only been mentioned in a doctoral dissertation (Dunlap, 1975). Their taxonomic frameworks of both the algal food and the animal were subsequently re-evaluated. Under current DNA-based taxonomy, we demonstrate that PoB is capable of feeding on multiple species of udoteacean algae. Feeding was observed under illuminated conditions, making it unlikely that PoB feeds exclusively at night. The ability to induce feeding under controlled conditions represents an important step toward establishing a culture system for PoB. Such a system is expected to provide a valuable experimental platform for elucidating the physiological, molecular, and evolutionary bases of kleptoplasty.

## Acknowledgements

The computation was performed using Research Center for Computational Science, Okazaki, Japan (Project: NIBB, 24-IMS-C346, 25-IMS-C302). We are grateful to Mr. Kazuki Nishida (Iwo World Kagoshima City Aquarium) and Atsushi J. Nagano (Nagoya University, Japan) for their help in keeping our collaborative work on track.

## Data Availability

All data and analysis codes are available at Figshare :10.6084/m9.figshare.31055917.

**Supplementary Table 1. Data of related algal species used for species identification**

**Supplementary Figure 1. Original maximum-likelihood phylogenetic tree based on *rbcL* sequences (715 aligned positions) of related Bryopsidalean algae**.

This tree served as the source topology for the phylogenetic representations shown in Figure 2

**Supplementary Video 1. Feeding of *Plakobranchus ocellatus* on *Rhipidosiphon lewmanomontiae***

**Supplementary Video 2. Feeding of *Plakobranchus ocellatus* on *Ventalia* sp. 4**

