## Supplementary figures and images for "Direct observation of feeding behavior in the kleptoplastic sea slug *Plakobranchus ocellatus* type black (Gastropoda, Sacoglossa)*"

### Supplemental Figure 1

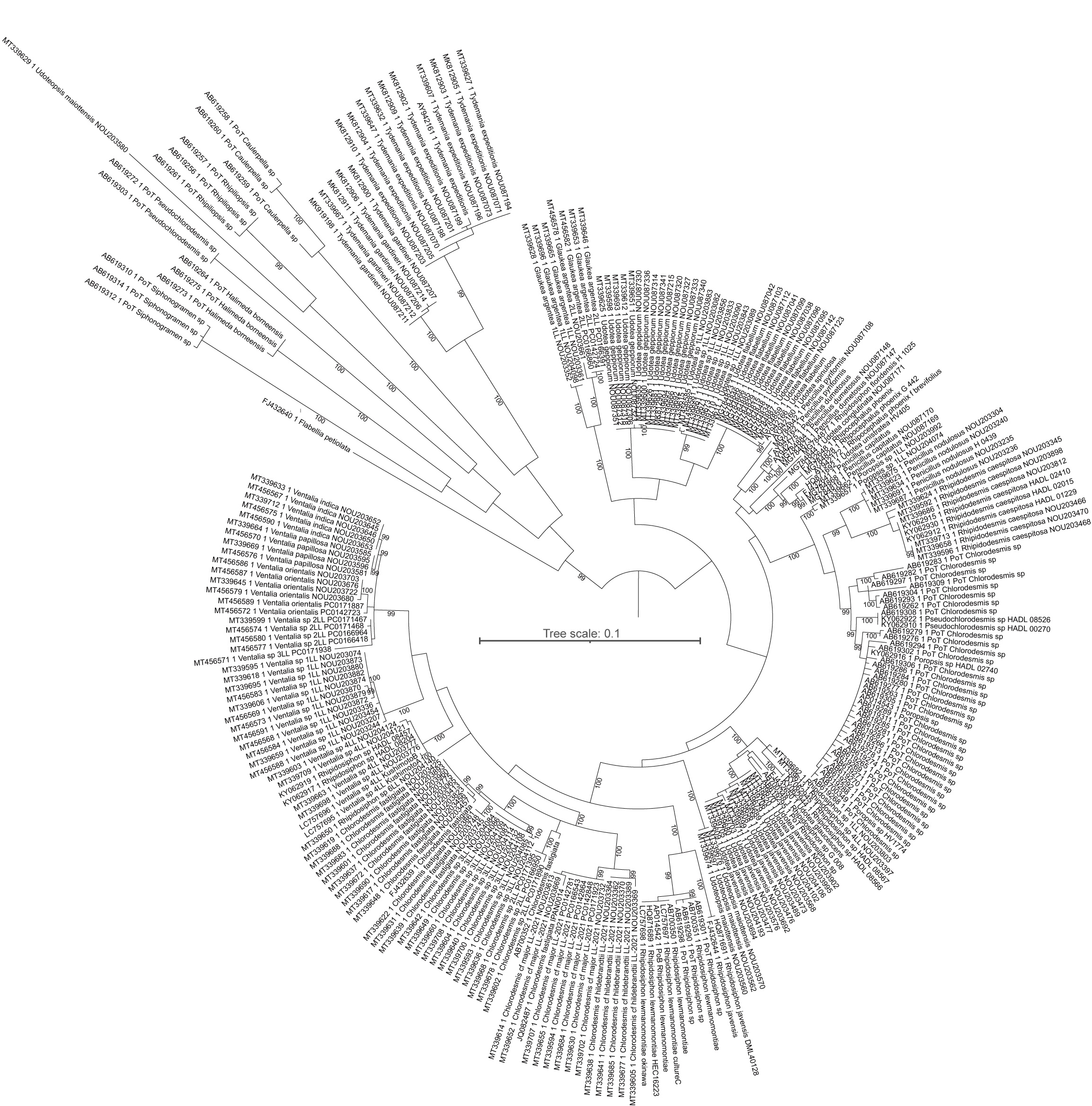
